# *De novo* Identification of DNA Modifications Enabled by Genome-Guided Nanopore Signal Processing

**DOI:** 10.1101/094672

**Authors:** Marcus Stoiber, Joshua Quick, Rob Egan, Ji Eun Lee, Susan Celniker, Robert K. Neely, Nicholas Loman, Len A Pennacchio, James Brown

## Abstract

Advances in nanopore sequencing technology have enabled investigation of the full catalogue of covalent DNA modifications. We present the first algorithm for the identification of modified nucleotides without the need for prior training data along with the open source software implementation, *nanoraw*. *Nanoraw* accurately assigns contiguous raw nanopore signal to genomic positions, enabling novel data visualization, and increasing power and accuracy for the discovery of covalently modified bases in native DNA. Ground truth case studies utilizing synthetically methylated DNA show the capacity to identify three distinct methylation marks, 4mC, 5mC, and 6mA, in seven distinct sequence contexts without any changes to the algorithm. We demonstrate quantitative reproducibility simultaneously identifying 5mC and 6mA in native E. coli across biological replicates processed in different labs. Finally we propose a pipeline for the comprehensive discovery of DNA modifications in any genome without *a priori* knowledge of their chemical identities.

## Introduction

DNA modifications are essential across the three kingdoms of life^1^, and are used by cells for defense, gene regulation, cell differentiation, and the transmission of regulatory programs across generations. A host of assays have been developed to detect specific modified nucleotides, including and especially 5mC and 6mA, which are widely deployed by prokaryotes and eukaryotes ^2–4^. Techniques exist to detect a diverse group of epigenetic modifications through the observation of DNA Pol II kinetics leveraging Single Molecule Real-Time sequencing (SMRT-seq) platform^5, 6^. In particular, pioneering work^7, 8^ demonstrated the capacity to identify DNA methylation marks via the comparison of native versus amplified DNA through supervised machine learning. The SMRT-seq platform provides observations of DNA modifications through analysis of polymerase dynamics, which leads to the current requirement of deep read coverage in order to identify particular DNA modifications^5, 8^.

Nanopore sequencing technology confers the opportunity to identify modified nucleotides through direct observations of single-molecules through monitoring electric current. Several pilot studies have demonstrated the feasibility of using nanopore-derived information to identify methylation marks in native DNA^9–11^. To date, such studies have used a highly processed form of the data generated by the nanopore platform. Further, no software packages have been developed to interrogate and visualize the raw data in a human-interpretable fashion. Here, we present software that implements visualization to enable direct exploration, and automated statistical procedures to discover DNA modifications of, in principle, any form, even when the chemical identity of the modification is not known *a priori*. That is, we utilize unsupervised, rather than supervised statistical learning. We demonstrate the efficacy of our approach for three known marks, 4mC, 5mC and 6mA, in an artificial “ground truth” setting, and also in a well-studied laboratory strain of *E. coli*.

The identification of modified nucleotides without training data (Figure 1) requires the (nanopore) sequencing of a native and matched amplified DNA sample (where amplification is employed to produce unmodified DNA). We developed the open source *nanoraw* software package (pypi.python.org/pypi/nanoraw version 0.4.2; code repository github.com/marcus1487/nanoraw and documentation nanoraw.readthedocs.io) for the processing and visualization of raw data. The re-squiggled (Methods) raw signal for both native and amplified samples is compared at each base genome-wide leading to the identification of consistently modified bases, with discriminative power to accurately detect known marks in *E. coli*, and also the potential identification of new signals of unknown origin. Several similar approaches have been previously described^6, 9–13^, but require prior training datasets not only for new types of modifications, but for the same modification in a new sequence context. Additionally, the reliance of model-based approaches on known training data sets does not allow for detection of modified and unmodified nucleotides in close proximity. Table 1 summarizes the central benefits of testing-based (*nanoraw*) versus model-based^12, 13^ modified nucleotide detection methods. In our view, testing-based modified nucleotide algorithms may soon enable the description of the entire collection of modified nucleotides in a genome.

**Figure 1.**
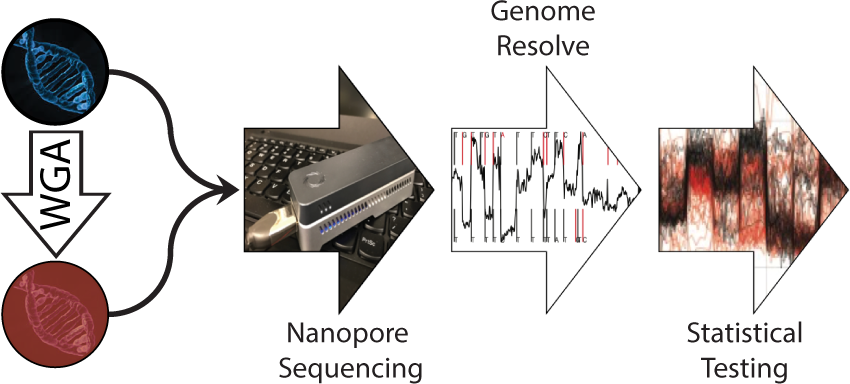
Modified DNA identification pipeline. Native and whole genome amplified (WGA) biological samples are processed using nanopore sequencing, raw signal is analyzed with nanoraw and statistical tests are performed to identify regions with modified bases.

**Table 1.** Testing-based versus Model-based Modified Nucleotide Detection

| Testing-based Advantages | Model-based Advantages |
| --- | --- |
| • No training data needed<br>• Identification of modified and unmodified nucleotides in close proximity<br>• Any chemical modification detectable | • Exact chemical modification known<br>• Exact modified position known<br>• Only native DNA needed after training |

We anticipate that this approach, enabled by the software we present, will play an important role in microbial, plant, and metazoan genomics, particularly and especially for non-model organisms. Many software packages exist^14, 15^ to assemble complete genomes from nanopore data, and hence genome sequence and epigenetic modifications can be simultaneously obtained in a single assay without prior knowledge of the collection of extant epigenetic marks in an organism. Further, we point to future work, where coupling to mass spectrometry and NMR may provide a complete parts list of endogenous DNA modifications in any system. The implications of this technology are clear and wide reaching for cancer genomics, population genetics, studies of epigenetic heritability, and the environmental biosciences.

## Results

### Visualization of the raw output of nanopore sequencing

Base-calling in nanopore sequencing currently relies on treating signal as a locally stationary process, first involving segmentation into stationary regimes (“events”), and then kmer-calling within segments to assign putative kmers^16–18^. Initial assignments are then resolved to individual nucleotide calls by joint analysis of consecutive segments. Precision for the initial k-mer calls is relatively poor, and is improved upon reconciliation of neighboring regions. Individual “1D” nucleotide calls show 83.9-89.6% identity in genomic alignments for single molecule reads (Table 2), facilitating both de novo genome assembly^15, 19, 20^ and the processing presented here.

**Table 2.**
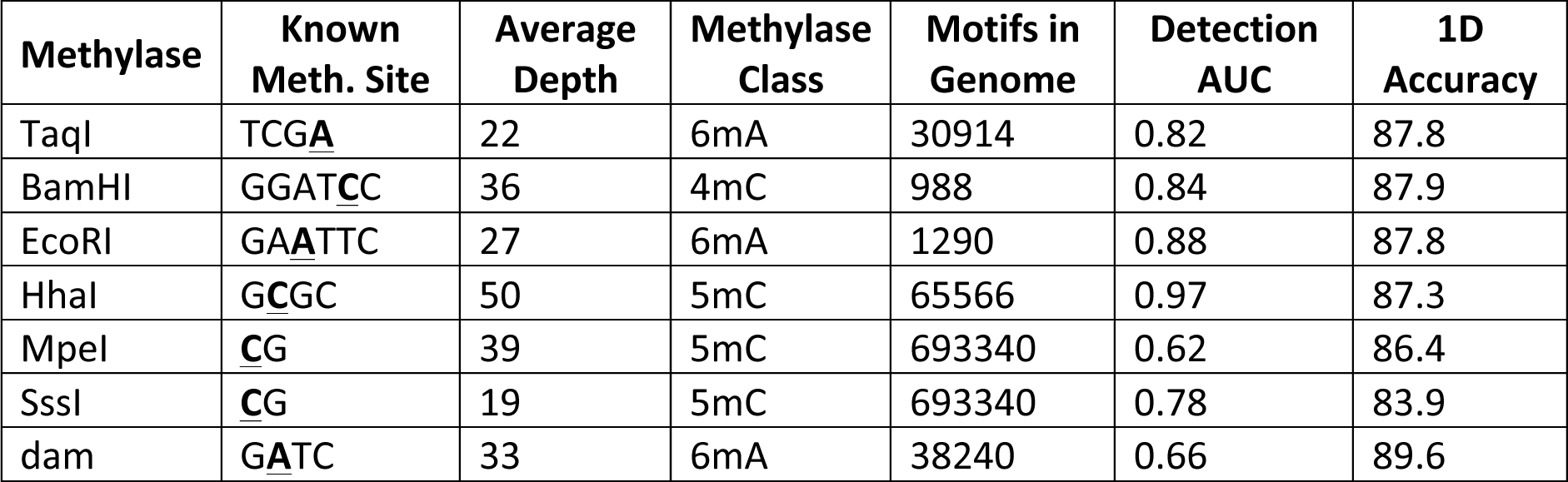
Tested methylases with known recognition site (methylated base underlined), depth of sequencing, methylation class, and other sample statistics.

We developed the *nanoraw* software package to precisely associate raw nanopore signal with genomic positions (Methods; implemented in the genome_resquiggle subcommand) and thus allow genome-browser style visualization of raw nanopore signal – a utility so far missing from nanopore software resources. The resolution of raw signal with aligned genomic sequence constitutes a robust procedure applicable to current and foreseeable subsequent generations of the technology with little to no tuning of parameters. The *nanoraw* software allows the selection of genomic locations via a multitude of criterion enabling the visual identification of regions of consistent or inconsistent raw signal and, as a result gains insight and intuition into the process of nucleotide assignments. Additionally, *nanoraw* provides a utility for outputting genome-wide statistics for groups of corrected reads in standard wiggle format (Supp. Figure 1; statistics include q/p-values, difference in signal between groups, read coverage, mean signal level, mean signal standard deviation, and mean event length).

### Identification of chemically modified nucleotides

We leverage the previously established^9, 11–13^ strategy comparing native to amplified DNA in the context of genome-guided analysis to discover modified nucleotides without *a priori* knowledge of their chemical composition or effect on the nanopore signal.

To demonstrate the feasibility of this approach for three major classes of DNA modifications, we constructed a ground truth dataset using seven purified methylases to introduce methylation to whole genome amplified *E. coli* DNA at known target sites (Table 2). These methylases catalyze the addition of a methyl group to the DNA to produce three distinct methylated bases: 4 methyl-cytosine (4mC; M.BamHI), 5 methyl-cytosine (5mC; M.HhaI, M.MpeI and M.SssI) and 6 methyl-adenine (6mA; M.TaqI, M.EcoRI and M.dam). Each of these samples and two control samples were processed by nanopore sequencing (Methods).

Individual reads, corresponding to single molecules, are aligned to a genome using extant methods^21, 22^. We use BWA-MEM for all mappings presented here, but make the graphmap algorithm available through *nanoraw* as an alternative option. We use only template (in sequencing) reads in the analysis presented here due to a known shift in complement read signal^12^, likely due to re-annealing of the DNA helix. Nanopore technology is likely moving away from 2D read technology, so this does not present any issues for future applications. We note that the *nanoraw* pipeline can easily be applied to a genome derived directly from the same nanopore data using established pipelines^14^ when analyzing organisms without a reference genome. Nanopore assemblies of *E. coli* have yielded accurate single-contig genomes (99.5% nucleotide identity,^23–25^). After alignment, raw signal is re-segmented using the *nanoraw* genome_resquiggle command to map contiguous raw signal to mapped genomic positions (Methods). Given two collections of corrected events, one corresponding to native DNA and another to amplified, the identification of DNA modifications is reduced to a statistical testing problem. This approach contrasts with previous DNA modification identification algorithms which model signal shifts and require new training datasets for each modification and genomic sequence context^9, 12, 13^. To identify genomic positions with shifted electric current, as compared to an amplified sample, the *nanoraw* pipeline employs the Mann-Whitney U-test^26^. As modified bases consistently shift the electric current at several bases surrounding the modified base (Figure 2A-C, Supp. Figure 2), Fisher’s method^27^ was applied across a moving window to produce final significance tests (Methods). Bases admitting statistically significant tests indicated regions with modified nucleotide(s).

**Figure 2.**
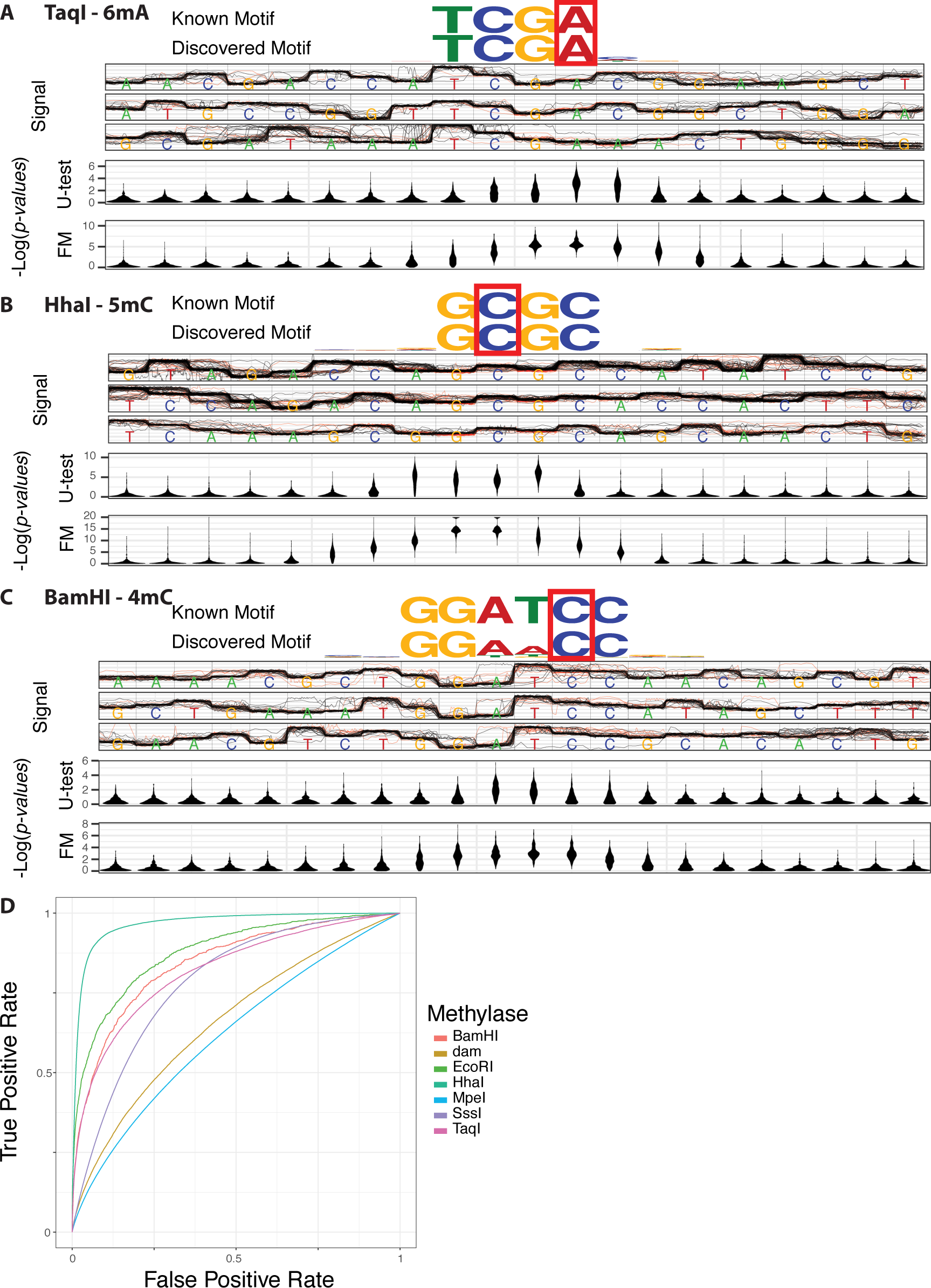
Identification of known modifications. **A.** Detection of three distinct chemical DNA modifications (4mC, 5mC and 6mA). Each panel contains the known methylase recognition sequence, the discovered sequence motif, the raw genome-anchored nanopore signal at the top identified regions (red – methylated reads, black - amplified reads) and distributions of Mann-Whiteny U-test and Fisher’s method negative log p-values at each base over the top 1000 most significant regions containing the motif. Additional tested methylases are shown in Supp. Figure 3 **B.** ROC curves for the seven methylase experiments show a range of capacity to discriminate known modification sites (determined by known motif).

For each chemical modification (4mC, 5mC and 6mA) *nanoraw* discovered the known sequence specificity of each enzyme based solely on shifted signal levels followed by a standard motif discovery pipeline (Figure 2, Supp. Figure 2; figures are immediately produced by the *nanoraw plot_motif_with_stats* command). Dam methylase shows expected motif degeneracy^5, 28^ and comparatively weak specificity. Mann-Whitney *p*-value (top of lower panel) and “smoothed” Fisher’s method p-value (bottom panel) distributions for the top 1,000 most significant regions containing the known motif (Figure 2, Supp. Figure 2, lower panel) indicate that globally the highest Fisher’s method significance values center on the known modified base. We note that the location of nucleotides with shifted signal is specific to the sequence context, as indicated by the U-test *p*-values. Borrowing strength across statistical tests at neighboring nucleotides (using Fisher’s method) yields a *p*-value distribution that generally achieves its minimum at the modified nucleotide, regardless of the sequence context. Additionally, six out of the seven methylases produce a lower AUC when testing either the nucleotide immediately up or downstream from the known modification (Table 2), indicating a preference for the peak of statistical significance to occur at the modified nucleotide.

Genome-wide, each methylase shows strong preference for the known sequence motif with variable levels of accuracy (Figure 2D; area under the curve (AUC) from 0.62 to 0.97; Table 2). For M.BamHI, the AUC can be improved from 0.84 to 0.93 by including the discovered degeneracy at the fourth position of the motif (Figure 2B), indicating potential preference for near-cognate sequences. Such star activity may be responsible for the reduced fidelity of detection of the known motif for several methylases tested here. Statistical power for the detection of modified bases also scales as expected with sequencing depth (Supp. Figure 3). Additionally, *in vitro* methylase activity may be lower for some methylases contributing to poorer detectability.

### Endogenous modifications in laboratory strain of *E.coli*

To simultaneously assess our capacity to identify endogenous modifications along with the biological and technical reproducibility of our approach, we applied our modified nucleotide detection pipeline to *E. coli* (strain K-12 MG1655), one of the best-studied genetic model organisms, independently in two laboratories (Experiment A: Pennacchio Lab, LBNL, USA, and Experiment B: Loman Lab, Univ. Birmingham, UK). Due to inclusion of two additional bacterial samples the coverage of the *E. coli* genome from experiment B samples was substantially lower (8X native and 6X amplified average strand-specific coverage) than the experiment A samples (98X native and 13X amplified average strand-specific coverage). In both experiments the expected^29, 30^ modifications catalyzed by M.dam (6mA) and M.dcm (5mC) were identified as the strongest *de novo* identified motifs (Figure 3A-B and Supp. Figure 4) and ubiquitously throughout the genome with similar specificity to *in vitro* methylation experiments (Figure 3C); Experiment A detects the M.dam and M.dcm targeted modifications with 0.85 and 0.93 AUC and experiment B with 0.73 and 0.90 AUC.

**Figure 3.**
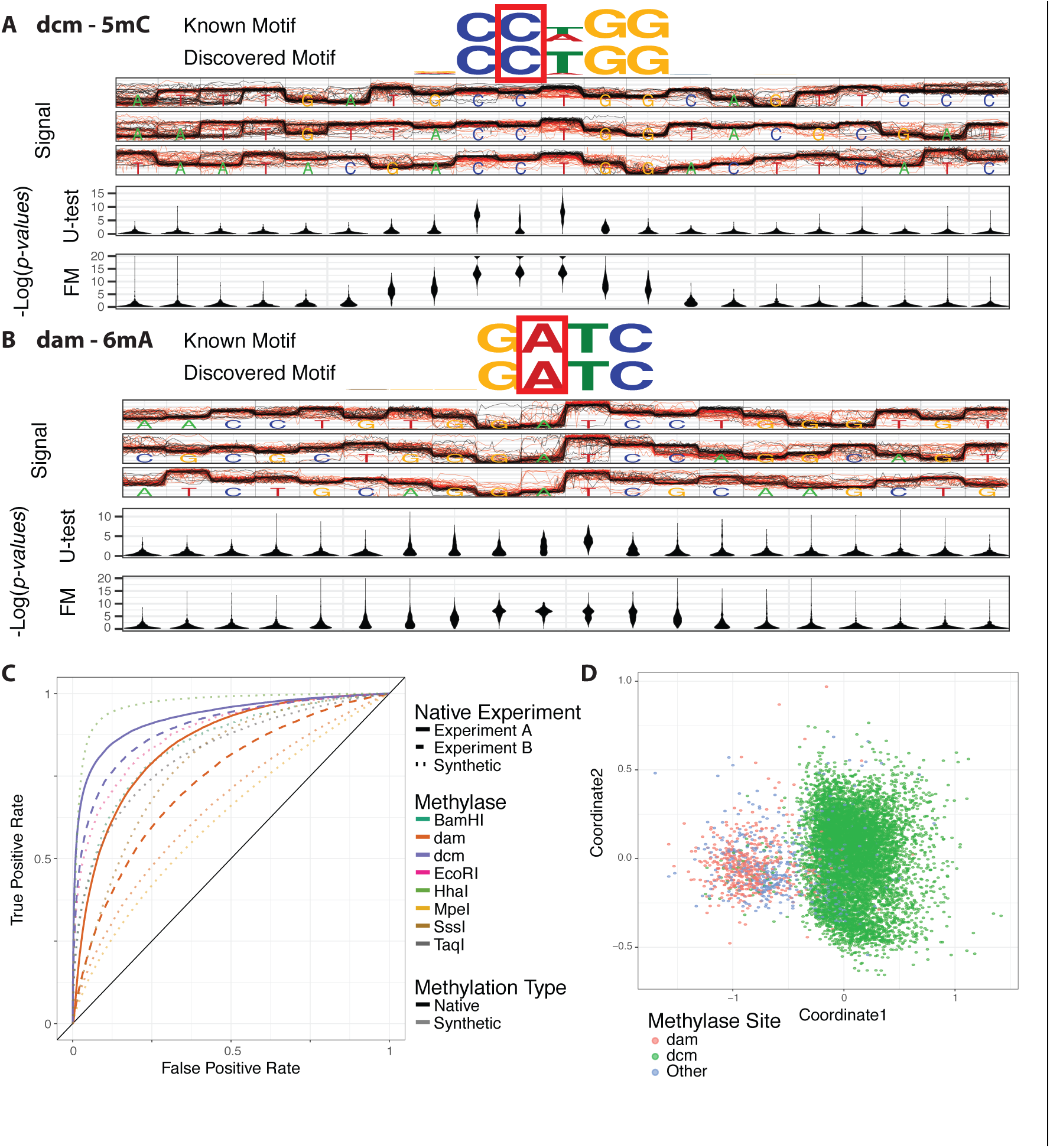
Identification of Native DNA Modifications. **A, B.** Two major classes of modifications found in native E. coli (dcm and dam methylase). Panels are as in Figure 2A. **C**. ROC curves for native modified bases as in Figure 2B. **D**. Clustering of individual modified locations based solely on raw genome-anchored signal (raw versus amplified). Color indicates motif the region matches.

In experiments A and B respectively, 97% and 88% of the top 1000 most significant sites are attributable to the known M.dam or M.dcm sequence motifs (Supp. Figure 6 and 7; within 2 base pairs). Indeed the top of the rank lists from the two experiments is greatly enriched for overlap; the very top of the rank lists are enriched 200 fold over random detection (Supp. Figure 8A). To assess the biological versus technical variability we down-sampled reads from experiment A and found that correspondence between pseudo-down-samples is approximately equal to the correspondence between equal depth samples from two experiments. We find that depth of sequencing is the driving factor for reproducibility in our study. Fraction of overlap between the top 1% of sites scales with coverage showing increased power to detect known motifs out to greater than 15X coverage (Supp. Figure 9). When only sites with greater than 11X coverage are considered we observe greater than 32% overlap between the two experiments within the top 1% (Supp. Figure 8). An extended discussion of factors affecting reproducibility of identification of DNA modifications can be found in the supplemental text.

### Modification Clustering

To discover relative signatures of modifications, we employed an unsupervised dimension reduction approach. For each identified (statistically significant) genomic position, we computed the difference between the native and amplified signal at each nucleotide within a window two bases up- and down-stream. This provides a signal shift signature for each modification/context (Methods). These signatures are compared using Euclidian distance (at allowed offsets up to 2 bases) and projected to 2-dimensions using Multidimensional Scaling (MDS) for visualization (Figure 3D). The two dominant clusters correspond the known 4mC and the 5mC methylases. Other sites contain the potential for modifications of unknown origin as well as potential false positives. Hence, unsurprisingly, the vast majority of epigenetic modifications in *E. coli* are of known origin, and our pipeline detects them and provides a clustering that segregates the underlying modification. This analysis demonstrates that the raw signal can be used to visualize, cluster, and detect distinct DNA modifications genome-wide.

### A note on signal normalization

In addition to resolving raw signal with a genomic alignment and appropriate statistical testing, raw signal normalization is key to the accurate identification of modified bases. Most current nanopore base calling and methylation detection algorithms ^12, 13, 16–18^ utilize picoamp (pA) values produced by normalization of the raw 16-bit data acquisition (DAC) values. This normalization occurs in steps, first taking the mean of DAC values over detected events, followed by application of instrument-derived parameters (offset, range and digitization) to produce raw pA estimates. These raw pA estimates contain systematic shifts in signal over each read (Supp. Figure 10). These values are then corrected by fitting of k-mer dependent shift and scale parameters ^12^. This fitting procedure requires that the read be basecalled such that a k-mer is assigned to each estimated event as well as a lookup table of standard k-mer current levels derived from many runs on the same nanopore chemistry. All normalization procedures (DAC, raw pA, pA and median) are made available in the *nanoraw* package.

To discover modification directly from raw signal, we apply median normalization (Methods) based solely on the raw DAC signal values (without use of segmentation, called bases or instrument-derived drift parameters). Despite the relative simplicity of the median normalization technique we see that both both pA and the far more straightforward median normalized signal explain 76% of variance for 4-mers (Supp. Figure 10; similarly for 6-mer we see 77% and 76% of explained variance for pA and median normalizations respectively). Note that the fraction of explained variance is invariant to scale, thus allowing comparison of normalizations on drastically different scales. We propose that median signal normalization constitutes a useful raw signal normalization method without the need for basecalling or the *a prior* generation of k-mer-level lookup tables, and that fraction of variance explained by k-mers constitutes a useful metric for the assessment of signal normalization procedures.

## Discussion

The methodology presented here allows for the identification of DNA modifications using raw nanopore data and statistical analysis without external or additional training data. At present, we require repeated observations of modifications at any genomic position for detection, meaning that individual modified nucleotides need to be consistently present at a genomic location in several independent reads, as with most current modified nucleotide detection assays (including Illumina-based assays such as bisulfite sequencing). Once identified, modeling of consistent signal shifts could conceivably generate phased, single-stranded modification maps in individual, single-molecule reads. As nanopore median and maximum read lengths continue to increase, these calls could be used to phase an epigenome with single-strand resolution. In diploid organisms, such as human, this would be a considerable advance over bisulfite sequencing and antibody-based approaches, and will likely soon enable the study of population epigenetics.

Iterating the base-calling procedure to explicitly take into account modifications during sequencing, progressively enlarging the chemical vocabulary of a base caller, will, at least in part, ameliorate the combinatorial complexity associated with this task. Indeed, one of the most obvious applications of *nanoraw* is the generation of training sets for base-calling algorithms that seek to improve the state of the art in this sequencing platform. Iteratively refined base-calls may improve resolution and power for the detection of modifications. Importantly, *nanoraw* does not require *de novo* knowledge of modification identity or sequence specificity. Thus training data sets can be produced wherein modified and unmodified bases exist within very short distances, as is the case in biological samples. Such training sets will likely prove useful to increase the accuracy of nanopore sequencing.

In organisms and systems with more diverse DNA modifications, or less sequence specificity at modified residues, our clustering approach provides an avenue for the systematic discovery of modified nucleotides, even when we do not know the identity of the modifications *a priori*. For instance, one could couple the strategies presented here to mass spectrometry (MS) or nuclear magnetic resonance (NMR) to discern specific moieties. One approach would be to fragment native DNA and use biotinylated oligonucleotide probes to pull down suspect regions of native DNA identified by *nanoraw* e.g. via sonication or the use of restriction enzymes, and then subject the precipitate to tandem mass spectrometry. In this way, a complete survey of DNA modifications could be derived *de novo* without the need for antibodies to specific moieties or enzymes. Given recent descriptions of the likely importance of 6mA modifications in metazoans, and the opportunistic (antibody enabled) nature of such discovers to date, it seems unlikely that a complete vocabulary of endogenous DNA modifications exists for complex organisms.

As the consistency of raw nanopore signal improves, it may become possible to identify modified bases from individual molecules without the need for repeated observations. This would open new furrows in exposure biology, where adducts are distributed stochastically in the genome due to non-endogenous chemical activities. More than 200 different types of DNA adduct resulting from exposure to exogenous and endogenous DNA binding compounds have been described^31^. These observations could be correlated with patterns of mutation in tumors and cell lineages within tumors to study the mechanisms of DNA repair underlying individual and environmentally-induced cancer susceptibility. Ultimately, such technologies may enable new diagnostic and therapeutic strategies in precision medicine.

Additionally, we acknowledge the enormous power of the human end-user for the detection of interesting patterns in complex data. The effective visualization of raw nanopore signal in genomic contexts may yield unexpected dividends as biologists browse signal-level information in regions containing genes or genomic elements of interest. For instance, we anticipate the generation of “ChIP-nano” assays to discover patterns of epigenetic marks associated with transcription factor binding sites. Such correspondences seem likely given a recent report that at least 70% of transcription factors in *Arabidopsis thaliana* have differential binding affinities at 5mC sites^32^.

Lastly, direct RNA sequencing will likely soon be possible on the Oxford Nanopore platform, and the approaches we present here may be useful for the study of RNA modifications. Combining the strategies presented here with a variety of pull-down assays for DNA and RNA has the potential to transform our understanding of the molecular codes of life.

## Methods

### DNA Sample Preparation and Nanopore Sequencing

Standard sample preparation including DNA extraction, whole genome amplification and nanopore library preparation (all samples presented here are “2D” reads) are described in detail in the supplementary methods. *In vitro* DNA methylation procedures are described in full in the supplemental methods.

### Resolve Indels Using Raw Nanopore Observations

Raw nanopore signal is processed first by segmentation into “events”, followed by assigning bases to those segments and joining of selected neighboring segments. This produces estimated base calls that contain errors. Individual reads are then aligned, using extant software, to a genome. We then resolve differences between the reads and the underlying genome sequence by re-segmenting the raw nanopore signal at genomic alignment insertions and deletions. Raw nanopore signal is anchored at stretches of alignments without indels. Resolving the segmentation of the raw signal to match the known bases constitutes the base algorithm used for all analyses presented in this manuscript. The full algorithm description can be found in supplemental methods and is implemented and publicly available in the open source python package *nanoraw* via the *genome_resquiggle* subcommand (pypi.python.org/pypi/nanoraw version 0.4.2; code repository github.com/marcus1487/nanoraw; and documentation nanoraw.readthedocs.io).

### Statistical Testing for the Identification of Modified Nucleotides

With raw nanopore signal assigned to each genomic base, comparison of raw signal levels between two samples is reduced to a testing problem. In order to test the difference between two samples the mean signal for each read at a genomic base are computed. We propose the Mann-Whitney U-test^26^ to test for differences in median signal intensity between two samples of interest. A robust order statistic-based approach is chosen as signal shifts near modified bases appear consistent, but not necessarily large in scale which other tests (e.g. t-test) have increased power to detect. The U-test is applied at every position across the genome with sufficient coverage (at least 5 reads in both samples). Since signal is affected at several bases surrounding a modified base, Fisher’s method^27^ for combining p-values is computed on a moving window (of two bases up and down stream for all tests in this paper) to produce final p-values.

### Signal Level Normalization

Nanopore sequencing is originally recorded as 16-bit integer data acquisition (DAC) values measuring the electric current across the nanopore. When the raw nanopore signal is converted to events (estimated base locations) Oxford Nanopore Technologies converts this integer signal value to an raw picoamp (pA) level via three parameters: offset, range and digitization. These pA values are then further corrected for pore and time specific drift of current level. This procedure requires a base called reads and a k-mer current lookup table. These normalization techniques are both implemented in the *nanoraw* package. Thus far, these measurements have been the gold standard for downstream analysis of nanopore signal levels.

To investigate and compare alternative normalization methods the following metric is proposed: fraction of conditional variance explained by k-mer (here we use 4-mer). We apply a median normalization procedure as the default for downstream signal processing. For a read with *N* raw signal observations (where *R*_*i*_ is the raw signal level at the *i*^*th*^ observation) we define the median normalized signal (*M*_*i*_) as *M*_*i*_ = (*R*_*i*_ − *median*_*j*∈[1,*N*]_(*R*_*j*_))/*MAD* where *MAD* = *median*_*i*∈[1,*N*]_(|*R*_*i*_ − *median*_*j*∈[1,*N*]_(*R*_*j*_)|). All signal measurements presented in this manuscript have been median normalized (with the one exception for the comparison to pA normalizations). In addition we winsorize the signal to clip aberrant spikes at plus and minus five MAD for all normalization types.

### Modification Based Sequence Motifs

To identify the sequence preference for the sites identified from a given modified versus amplified experiment we first identify the top 1,000 unique genomic locations based on computed Fisher’s method p-values. Seven bases of context are included up and downstream around each identified location. The meme algorithm^33^ is then applied to these sequences in the ZOOPS model. The top hit based on value is reported.

### Clustering

Each region centered on a base with significantly deviated current (based on U-test p-value) is identified. Each region is identified by the differences between the native and amplified signal levels. The distance between each pair of identified regions is computed as the minimum Euclidian distance over a 5-vector allowing a shift up or downstream by 2 bases (as peak significance is not exact). The dimension reduction algorithm MDS is applied in order to visualize clustering of the data. This algorithm is available via the *cluster_most_significant* subcommand in *nanoraw*.

## Acknowledgments

We thank Kenneth H. Wan, LBNL, for helpful comments during manuscript preparation. Research was conducted at the E.O. Lawrence Berkeley National Laboratory and performed under Department of Energy Contract DE-AC02-05CH11231, University of California. RN would like to acknowledge the EPSRC (EP/N020901/1) and the European Union’s Horizon 2020 research and innovation program under grant agreement No 634890, “BeyondSeq”. We thank Jared Simpson for personal communications regarding signal normalization.

## Author Contributions

MS organized the project. MS conceived the analysis procedures with input from JB. MS developed and implemented the *nanoraw* software package and nanopore analysis pipelines. MS, RE and JB reviewed processing pipelines. RN completed *in vitro* methylation reactions. JQ prepared samples and libraries for all Univ. of Birmingham experiments. JL prepared samples and libraries for all LBNL experiments. NL managed work and study design at the Univ. Birmingham, and LP managed work and study design at LBNL. MS and JB prepared the manuscript with input from all authors. All authors reviewed and edited the manuscript.

## Author Information

J.Q. and N.J.L. have received travel expenses and accommodation from Oxford Nanopore to speak at organized symposia. J.Q. and N.J. L. have received an honorarium payment to speak at an Oxford Nanopore meeting. N.J.L. is a member of the Oxford Nanopore MinION Access Programme and has received reagents free of charge as part of the MinION Access Programme and in support of this project but does not receive other financial compensation or hold shares.

## Supplemental Material

### Supplemental Text

#### Overlap and reproducibility of identified modifications between replicates

Measuring and quantifying reproducibility of assays which identify modified nucleotides is inherently challenging. Unlike many biological assays where the top hits consistently show effect sizes much larger than the majority of other sites (such as ChIP-seq), all modified sites throughout the genome are essentially equally statistically powered (modulo a few factors). This means that if for example 50,000 sites within the *E. coli* genome are truly modified in 100% of tested DNA fragments then these 50,000 will be randomly re-ordered in terms of any statistical test from one replicate to the next. Additionally, the null distribution contains all other sites in the genome. For the relatively small *E. coli* genome this elicits ~8.5M strand specific tested sites. Under the null distribution p-values will randomly distribute even between 0 and 1. Thus by random chance, we would expect to see within the *E. coli* genome at least one non-modified base with a p-value as low as 10e-7. As the statistical test for the identification of truly modified sites is likely not powered to to this level, the 50,000 truly modified sites will be mixed within these randomly identified sites from the null distribution. As shown in the main text and Supp. Figures 6 and 8 strand-specific coverage has a strong effect on the reproducibility as increased coverage increases the statistical power.

In addition to these factors, methylation is well documented to change with cell growth phase^34^. Given that no attempt was made to synchronize the growth phase in either lab, this and other biological effects may contribute to the discovery of modified sites unique to one of the two experiments. Using a mixture model approach (Methods) to account for differential coverage, we estimate that genome-wide 96% of dam and dcm recognition sites are methylated within experiment A data while only 83% of such sites are methylated within experiment B (Supp. Figure 7). Globally we estimate 25% of sites within experiment A sample show a shift in signal (indicating that they are in close proximity to a modified nucleotide) as compared to 7% of sites from experiment B (Supp. Figure 7).

Finally, given that the nanoraw MoD-seq pipeline does not have single base precision in the current implementation, sites within several bases of a modified base may all obtain significant p-values. These sites contribute additionally to a lack of overlapping significant sites between two replicates.

### Supplemental Methods

#### Experiment Library Preparation (LBNL)

Total genomic DNA from *E. coli* str. K-12 substr. MG1655 was extracted using previously described methods^35^. In brief, DNA was extracted from approximately 4 × 10^9^ log-phase cells using the QIAGEN Genomic-tip 20/G according to the manufacturer’s instructions (Qiagen, Valencia, California). Whole genome amplification of *E. coli* total genomic DNA was performed using the QIAGEN REPLI-g Single Cell Kit according to the manufacturer’s instructions (Qiagen). DNA was quantified using Qubit dsDNA BR assay (Life Technologies, Grand Island, New York).

2D sequencing libraries were prepared from native and amplified E. coli DNA according to the ONT recommended protocol (SQK-NSK007). In summary, DNA was fragmented using a Covaris g-TUBE (Covaris, Ltd., Brighton, United Kingdom). The fragmented DNA then underwent DNA damage repair using the FFPE DNA Damage Repair Kit (NEB, Ipswich, Massachusetts) and AMPure XP bead clean-up (Beckman Coulter, Brea, California). The DNA was end-repaired and A-tailed using the NEBNext Ultra II End Prep Kit (NEB). Following AMPure XP bead clean-up, adapters were ligated onto the DNA using Blunt/TA Ligase Master Mix (NEB). Libraries underwent a clean-up step using MyOne C1 Streptavidin beads (Life Technologies) and were quantified using Qubit dsDNA HS assay (Life Technologies). All sequencing runs were performed using R9 flow cells and MinION Mk1b devices with the standard MinKNOW 48-hour sequencing protocol. Metrichor was used to perform basecalling using the 2D Basecalling for FLO-MIN105 250bps workflow.

#### Experiment Library Preparation (University of Birmingham)

Total genomic DNA was isolated from three bacterial cell pellets (*S. aureus, M. smegmatis* and *E. coli* K-12) using the genomic buffer set and 500/G genomic tips (Qiagen) following the manufacturer’s instructions and mixed in equal amounts. DNA was fragmented using a Covaris g-TUBE in a centrifuge at 5000 rpm. Part of the material was end-repaired and A-tailed using the NEBNext Ultra II End Prep Kit. Following AMPure XP bead clean-up, PCR adapters provided in the SQK-NSK007 kit (ONT) were ligated onto the fragments using Blunt/TA Ligase Master Mix (NEB). 10 ng of the cleaned-up, adapted fragments were PCR amplified using LongAmp Taq 2x Master Mix (NEB) and the primers provided in the SQK-NSK007 kit. Following 18 cycles of PCR fragments were cleaned-up and sequencing libraries were prepared for both PCR amplified and the native DNA set aside earlier according to the ONT recommended protocol (SQK-NSK007) described above. Sequencing runs were performed using R9 flow cells and MinION Mk1b devices with the standard MinKNOW 48-hour sequencing protocol. Metrichor was used to perform basecalling using the 2D Basecalling for FLO-MIN105 250bps workflow.

PCR amplified DNA that underwent methylase treatment were barcoded using the native barcoding kit (EXP-NBD002) so multiple treatments could be multiplexed on one flowcell. Approximately 200 ng input DNA for each treatment was barcoded and pooled according to the 2D Native barcoding genomic DNA protocol. A library was prepared from the pooled, barcoded fragments using the SQK-LSK208 kit according according to the ONT recommended protocol. Two sequencing runs were performed using R9.4 flow cells and MinION Mk1b devices with the standard MinKNOW 48-hour sequencing protocol. Metrichor was used to perform basecalling using the 2D Basecalling for FLO-MIN106 250bps workflow.

#### Synthetic DNA Modification

DNA methyltransferases were purchased from New England Biolabs and used according to the manufacturer’s instructions. The exceptions are the M.MpeI (Chrometra, Belgium) and M.TaqI, which was expressed and purified by the Protein Expression Facility, Birmingham. DNA methylation was performed in vitro by incubating DNA (60 ng/uL) with 1 uL of methyltransferase and 80uM S-adenosyl-L-methionine in 50 uL of aqueous solution containing the appropriate methyltransferase buffer (NEB Cutsmart Buffer was used for both the M.MpeI and M.TaqI). Reactions were incubated at 37C for 1h (60C for 1h for M.TaqI) and then purified directly for sequencing using SPRI magnetic beads.

#### Resolve Indels Using Raw Nanopore Observations

Raw nanopore data produced by the Oxford Nanopore Technologies MinoION device is stored as a digital integer value that represent a measure of electric current as DNA passed through a nanopore (at a current rate of 4000 observations per second). As DNA passes through a nanopore this signal changes as some function of the local base pair composition of that DNA molecule. For DNA this function has been resolved with considerable accuracy by Oxford Nanopore technologies, but significant errors remain in the 1D base calls (between 70%-90% accuracy reported though this depends strongly on the version of pore used^20, 21, 36^). These errors can make it difficult to process or interpret the signal associated with a particular position of interest on the genome, as is common practice in genomic sciences. Thus a key step to more exact and confident interpretation is to resolve base calls made from this raw nanopore signal with a known or discovered consensus genome.

In order to address this problem the following algorithm is proposed and implemented in the *nanoraw* software package to assign contiguous segments of raw nanopore signal with genomic positions. Starting from the Oxford Nanopore Technologies base calls the first step is to align base calls to a provided genome (this genome could even have been discovered and assembled *de novo* from the same run^14, 15, 19^). Currently, *nanoraw* allows the use of either graphmap^21^ or BWA-MEM^22^ long read mapping algorithms, but any long read aligner could be used. All alignments presented in this manuscript were completed with the BWA-MEM algorithm. Then stretches of correctly mapped regions (including matching and mismatched base pairs) are used to anchor the called nanopore segments to genomic bases. Then aligned insertion and deletions (indels) must be resolved to assign raw nanopore signal to the assigned genomic bases. We note that indels are extended to the smallest non-ambiguous region where a called indel could identically be represented by another pairwise alignment.

For insertions into the genome, there are segments of the raw nanopore signal that are assigned to base(s) that do not exist in the genome. When such a region is encountered the region is extended out to the neighboring segments and one new segment is determined from the raw signal (using the process described below). Conversely for deletions there are genomic bases that have no assigned signal. The region defined by events surrounding these deleted base calls are identified and the number of deleted base pairs plus one (for the two correctly aligned neighboring bases) segments are then identified from the raw signal in this region.

For both insertions and deletions the final stage is to identify a specified number of new segments within a stretch of raw signal. In order to accomplish this the running difference between the mean signals of neighboring regions (currently using 4 observation windows) are computed. The site within the region of interest with the largest difference in signal level is called as the first segment. Then the next highest peak site is chosen unless it is within four observations of a previously added segmentation site. This process is repeated until the requested number of segments is identified. It is possible that the requested number of segments cannot be identified, and in this case the neighboring correctly called segments are included into the region of interest and the processes is repeated to identify two more segments from the expanded region. If extending this indel region intersects another indel these two indel regions are merged and re-segmented together. For all reads presented in this manuscript if a read requires the re-segmentation of greater than 100 contiguous bases it is excluded from further analysis.

This algorithm is implemented in the *genome_resquiggle* subcommand of the *nanoraw* software package.

#### Identify Regions of Interest

In addition to the Mann-Whitney U-test, the t-test is a supported test in the *nanoraw* software for convenience though we found better identification of known methylated site with the robust Mann-Whitney U-test. Additional regions of interest that are query-able by the *nanoraw* software are regions of maximal coverage, regions centered on a k-mer of interest (e.g. homopolymers have proven to be difficult to process in nanopore data^16, 37^), and regions with the largest raw difference in signal means between two groups of samples. The genomic sequence at the most significant regions of interest are also available directly in fasta format via the *write_most_significant_fasta* subcommand and standard wiggle files are produced by the *write_wiggles* subcommand for genome-wide investigation. This collection of region identification tools gives the nanopore investigator incredible power to interpret and further develop the potential for this technology.

#### Filter to identify reads with most consistent and biologically relevant signal

In order to remove reads that appear to be of low quality we have developed a filter based on the number of observations per base (event). Given the resolution of the raw signal to the genomic alignment, we have much more accurate picture of how many observations are made per genomic base. Bases that contain many more observations indicate a “stuck” base. Sometimes this may be of biological interest, but signal level variance analysis indicate that the majority of reads with many “stuck” bases do not provide signal levels matching the trends for reads that pass through the pore at a consistently fast rate. We recommend a filter to remove any reads with greater than 5,000 observations at a single base or greater than 200 observations in more than 1% of the bases within a read and this filter is applied to all analyses presented in this paper.

#### Mixture Model Percent Modified Bases Estimation

In order to estimate the fraction of positions with signal affected by a DNA modification we employ a mixture model implemented in the R package fdrtools^38, 39^. This model attempts model a distribution of p-values with a uniform component (which represents the false tests) and a monotonically increasing component (representing the true positive tests). Here we report one minus the estimated fraction composed within the null (uniform) component as the fraction of sites affected by modified bases.

### Supplemental Figures

**Supplemental Figure 1.**
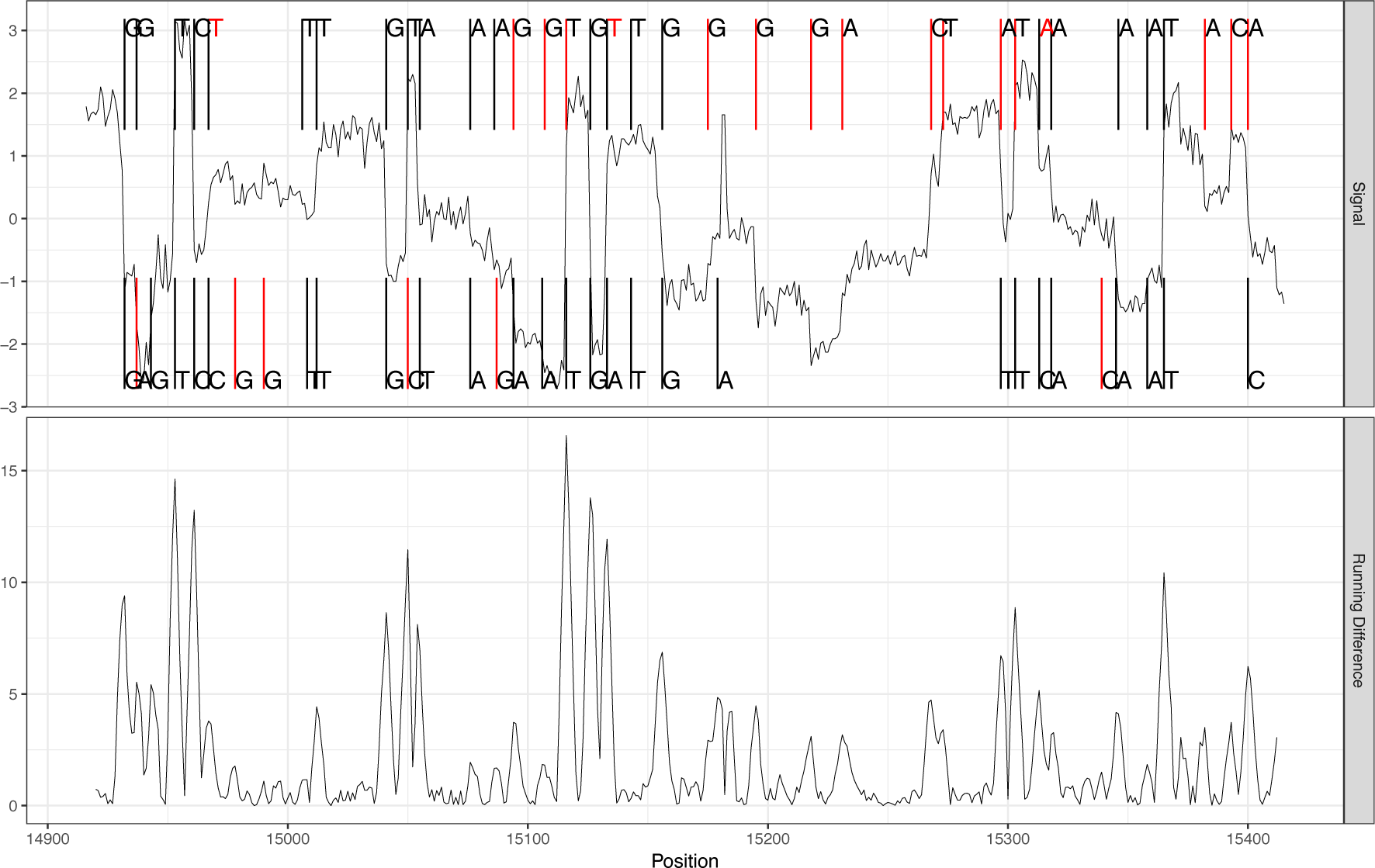
Example of corrected region of signal process is depicted. In the Upper panel segments on the top are estimated basecalls, segments on the bottom are genomic bases from a pairwise alignment and the signal through the graph is the median normalized raw nanopore signal (observed at 4kHz). Base call segments (top) marked in red indicate inserte bases, genomic segments (bottom) in red indicate deleted bases and called bases marked in red indicate mismatches. The lower panel shows the moving window of differences between neighboring windows of 4 observations. This measure is used to identify new segments.

**Supplemental Figure 2.**
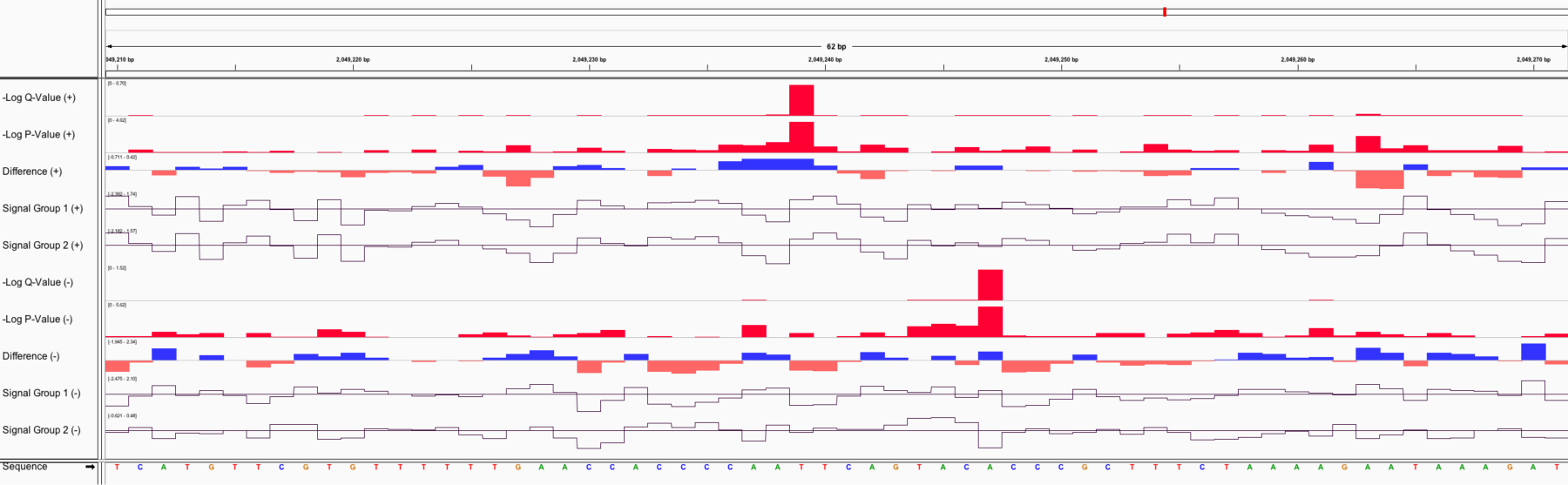
Example genome browser locus showing statistics for methylated and un-methylated genome resolved nanopore data. Showing (from top to bottom; second set of tracks are reverse strand) q-value, p-value, difference in signal between groups, mean signal from group 1 (un-methylated), and mean signal from group 2 (methylated). Not shown are coverage, mean signal standard deviation, and mean event length.

**Supplemental Figure 3.**
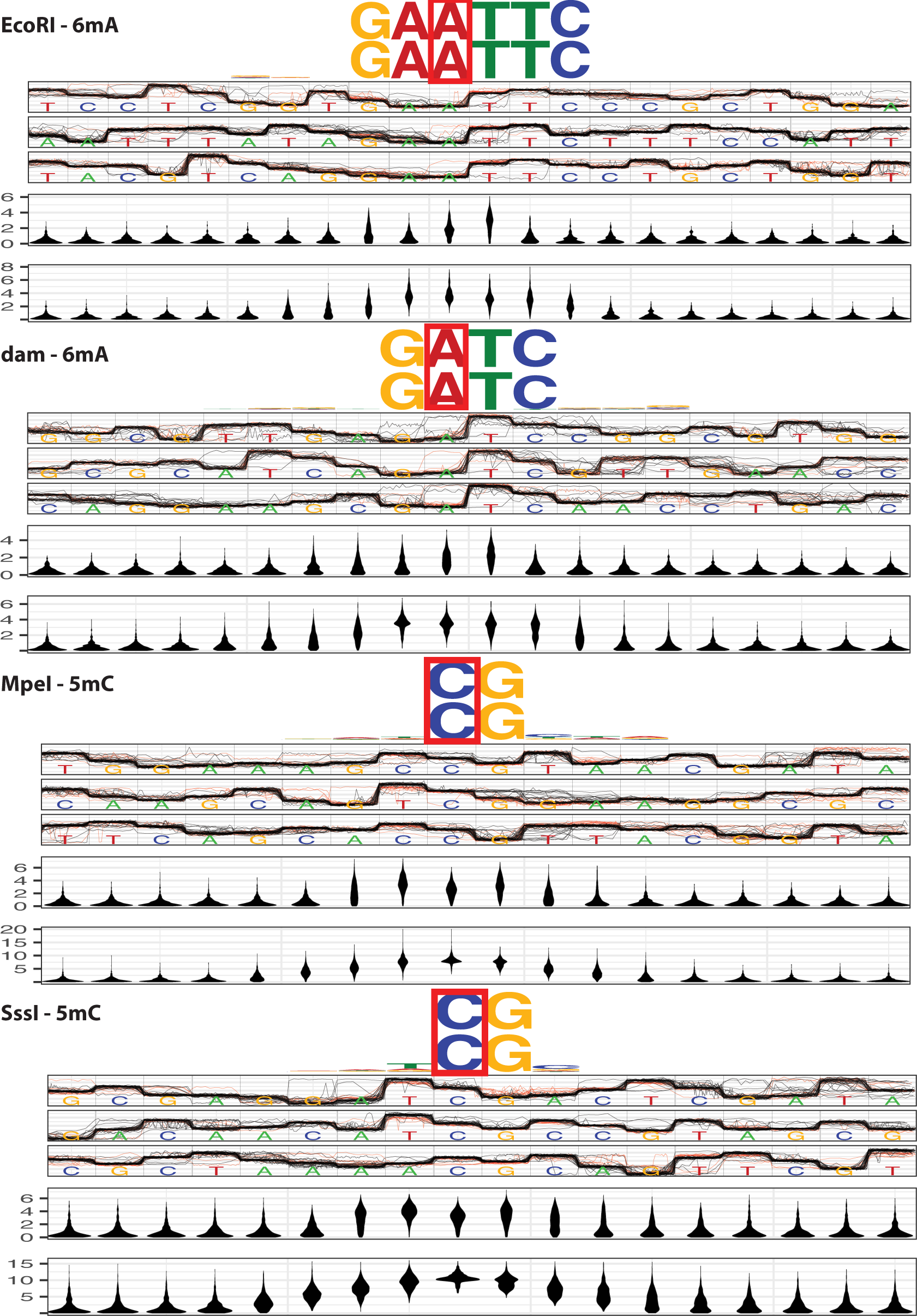
Additional known methylase signal plots (as in Figure 3). Red boxes indicate known methylation site.

**Supplemental Figure 4.**
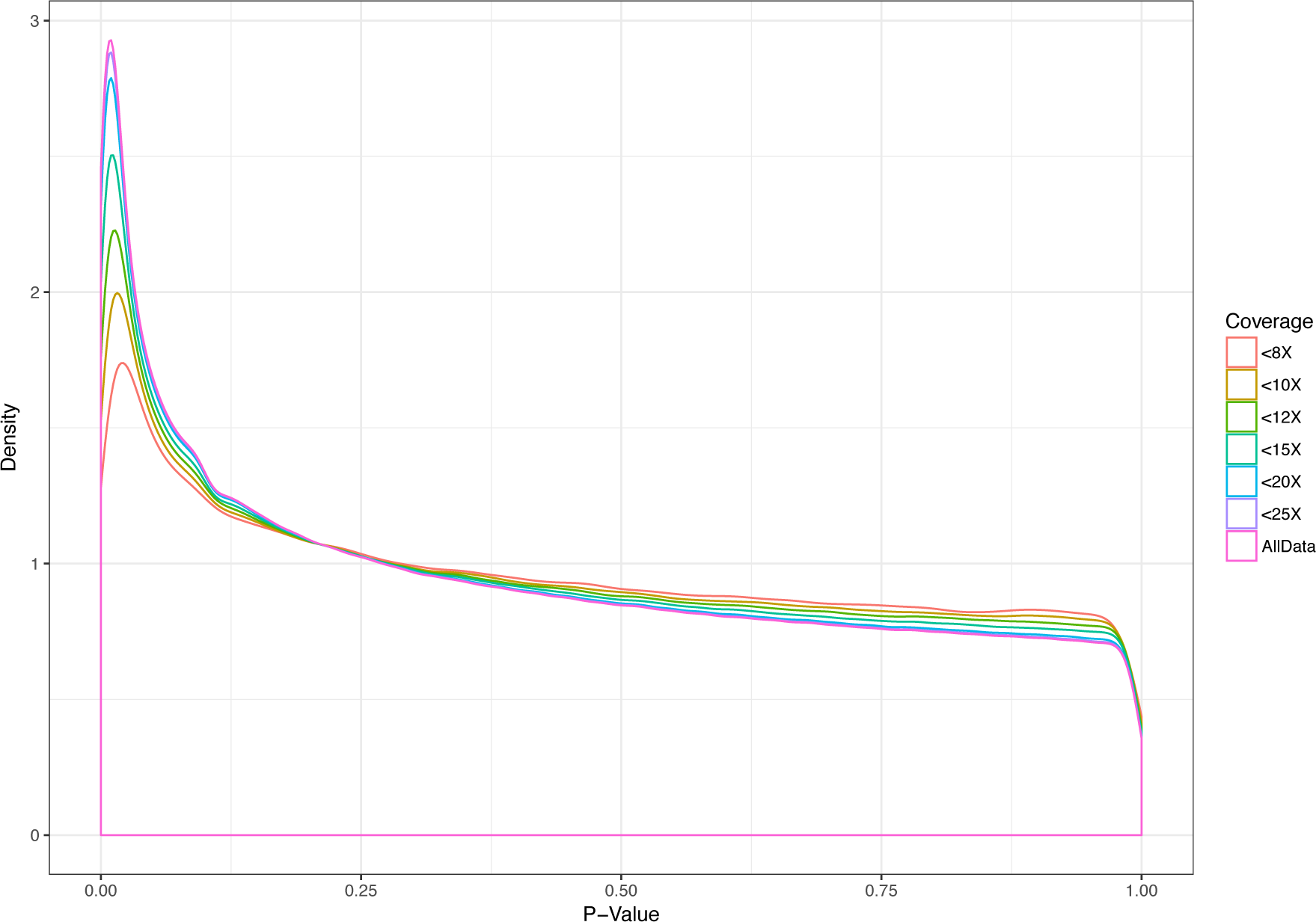
Distribution of p-values at given thresholds of minimum (between native and amplified) coverage at a site. Experiment A used for this analysis.

**Supplemental Figure 5.**
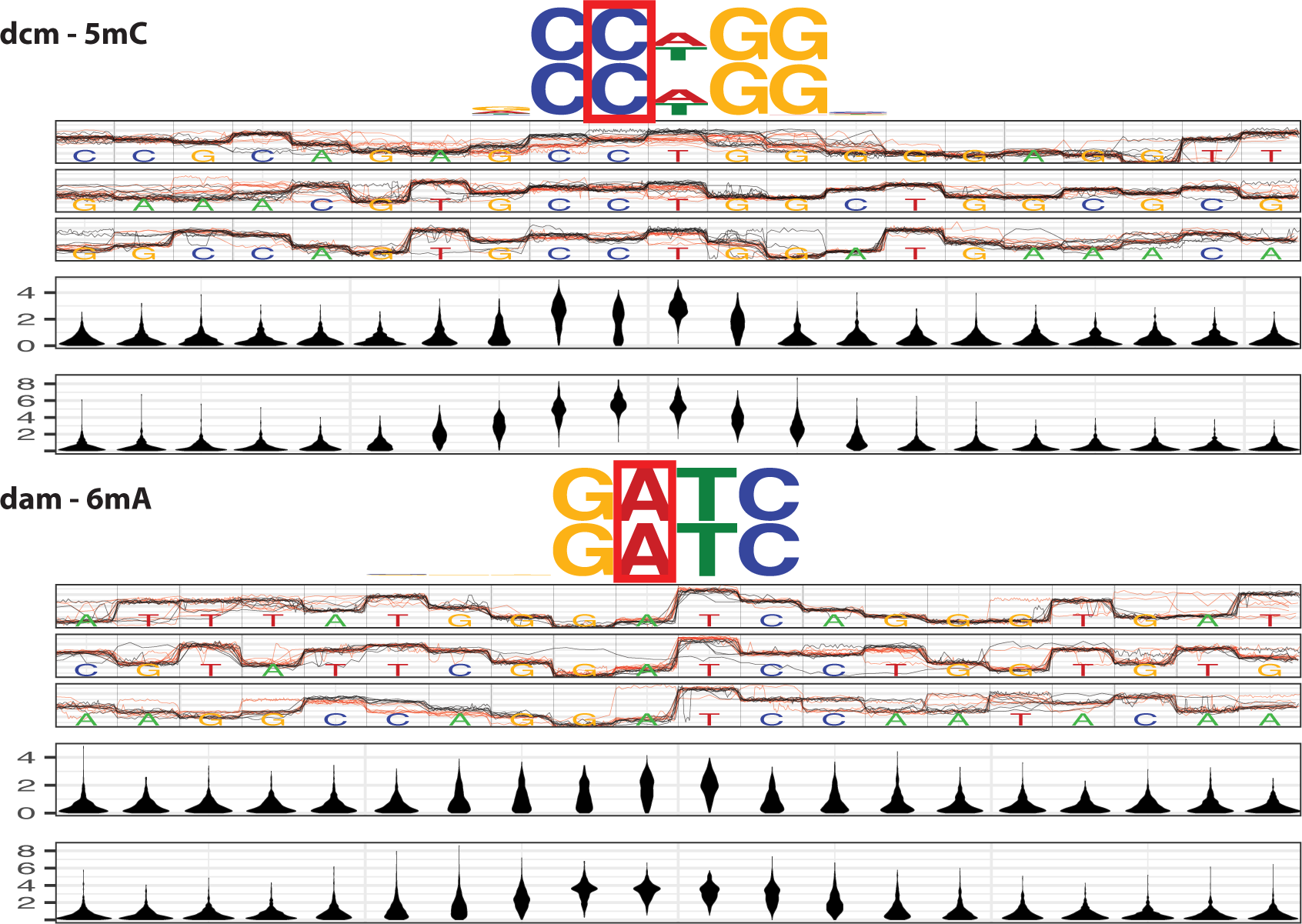
Examples for two major classes of modifications from lower coverage data from experiment B as well as negative log U-test and Fishers p-value distributions for 1,000 most significant sites within the known motif (as in Figure 4A). Arrows indicate known methylation site.

**Supplemental Figure 6.**
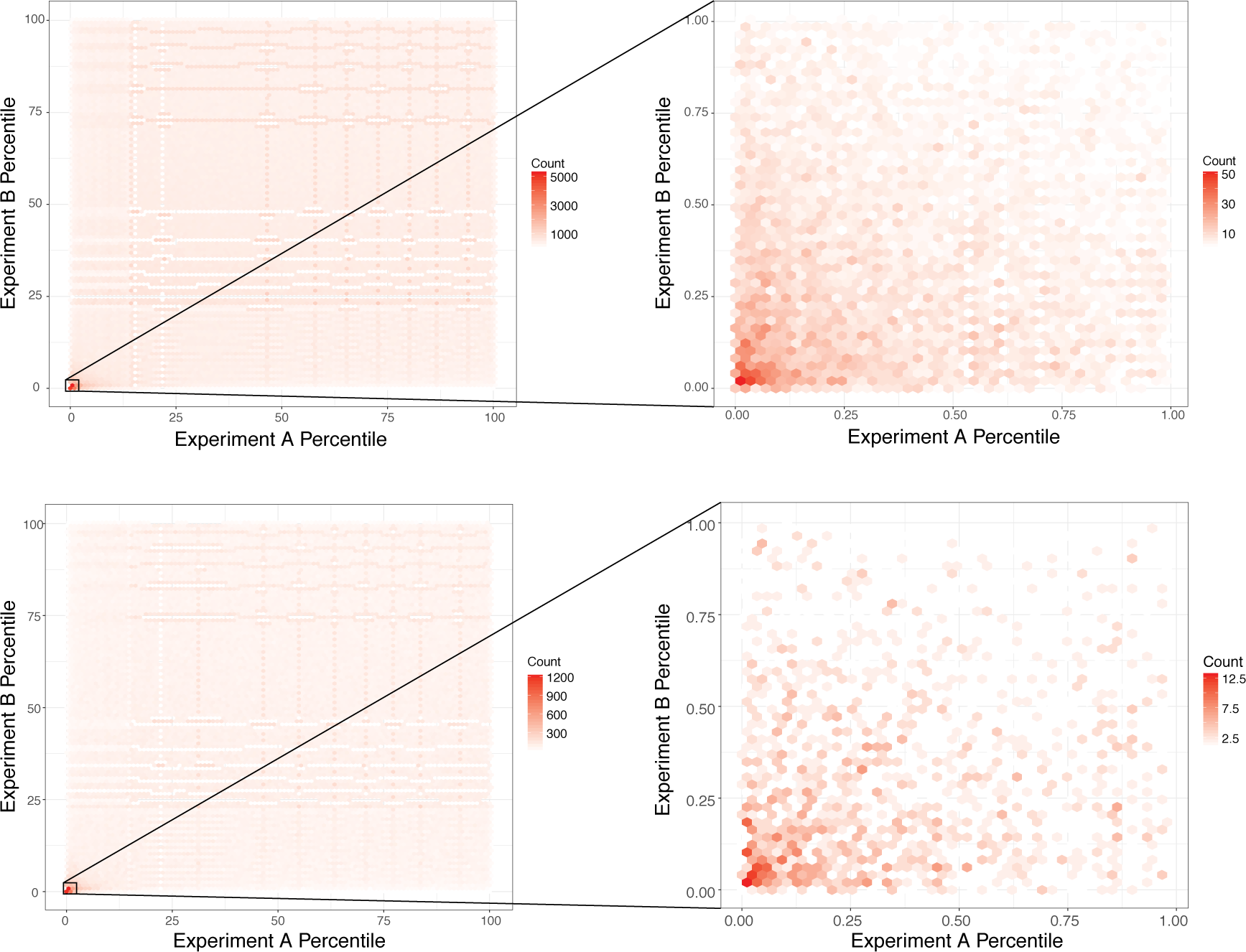
Density of sites showing correspondence between two experiments across rank lists (by p-value). Right panel is zoomed in to the top 1% of both lists. Lower panel shows the same correspondence restricted to sites with greater than 7X coverage (as opposed to 5X).

**Supplemental Figure 7.**
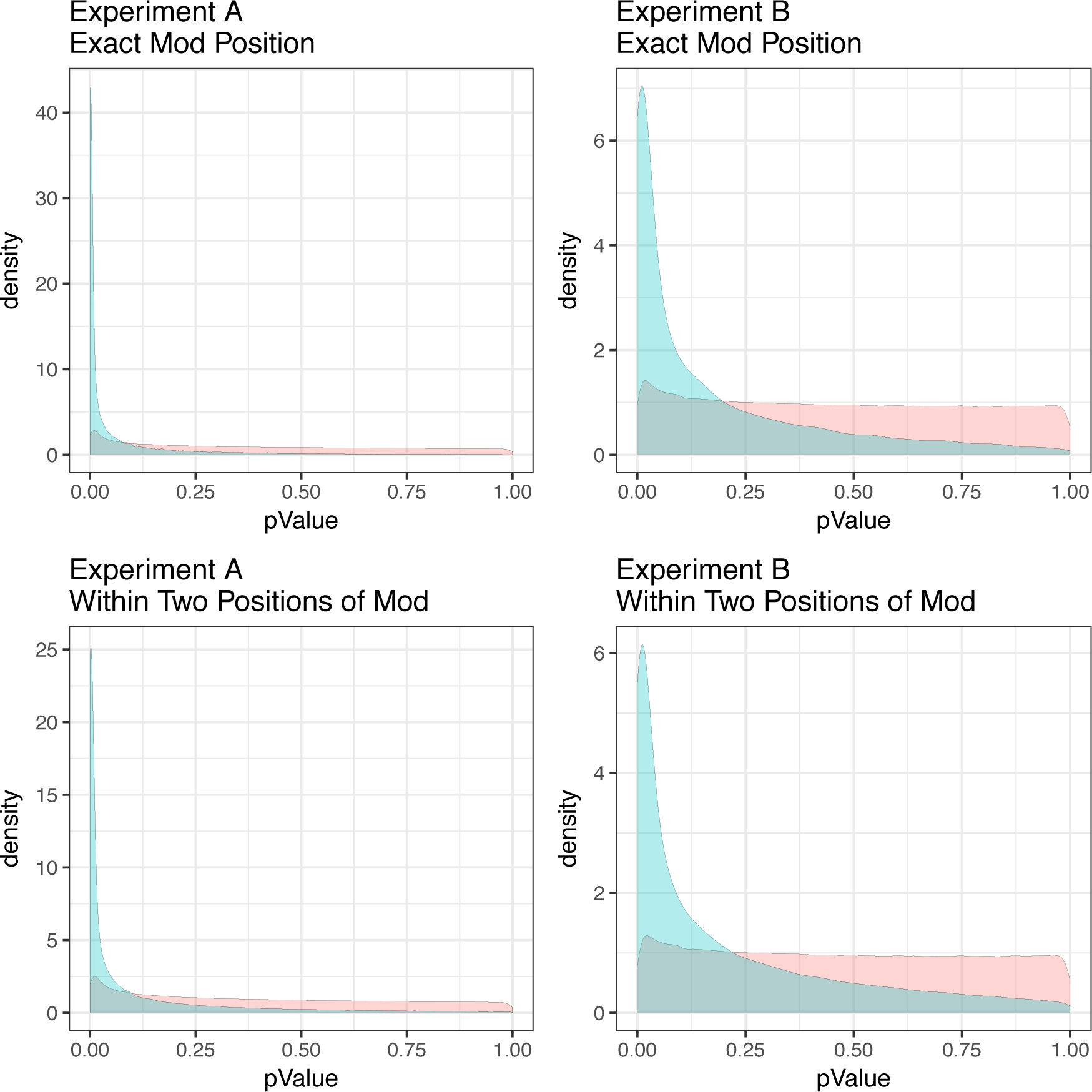
P-value distributions for experiment A (left) and B (right) across both regions that contain either dam or dcm motif and those regions that do not contain a motif.

**Supplemental Figure 8.**
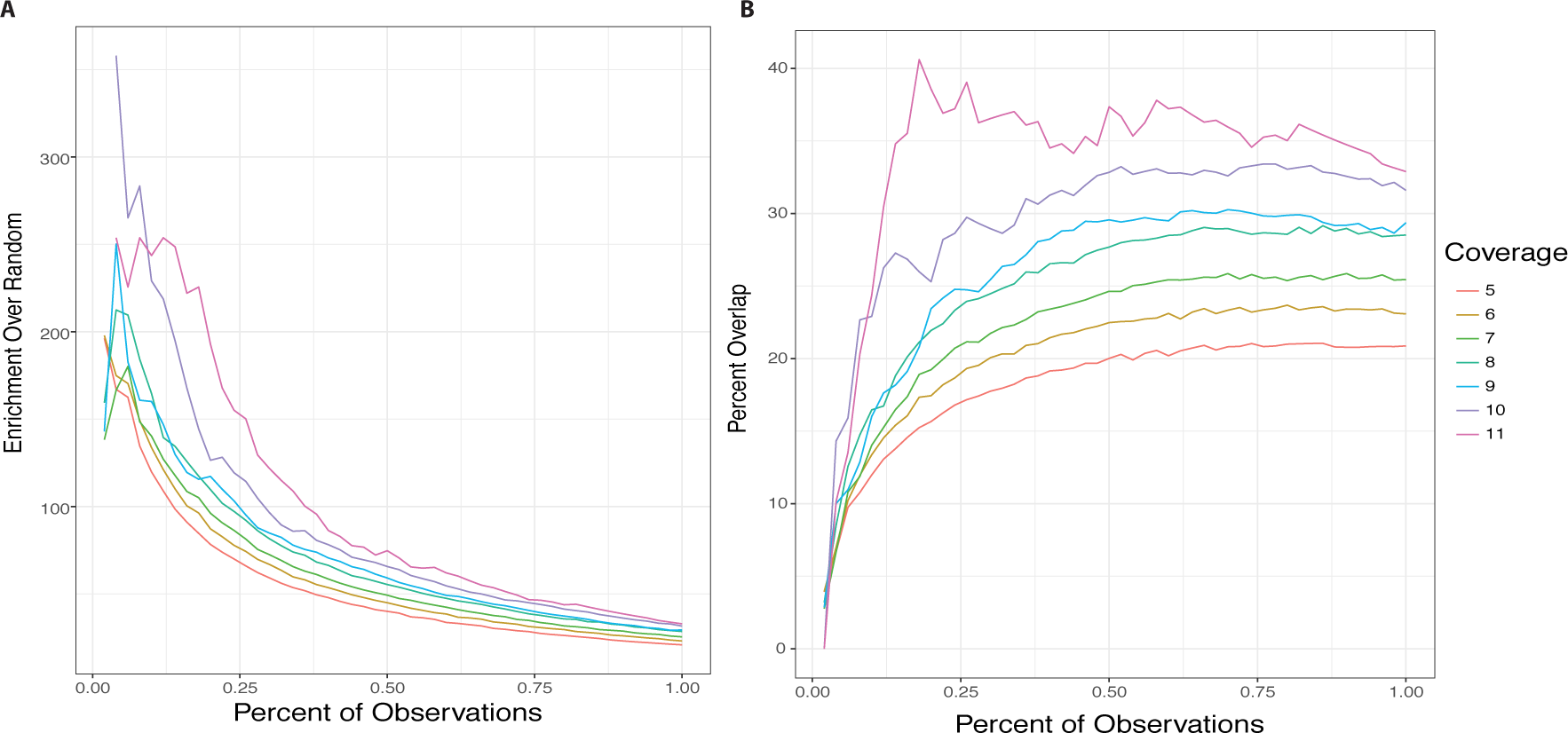
Comparison of modified bases identified by entirely independent processing pipelines including labs, technicians and sources. Moving down the rank list from both experiments (x axis) the enrichment over random (left panel) and percentage of overlap (right panel) is computed (y axis). Different lines indicate the minimal coverage filter applied to test each base across all four sequencing experiments (native and amplified from both experiments).

**Supplemental Figure 9.**
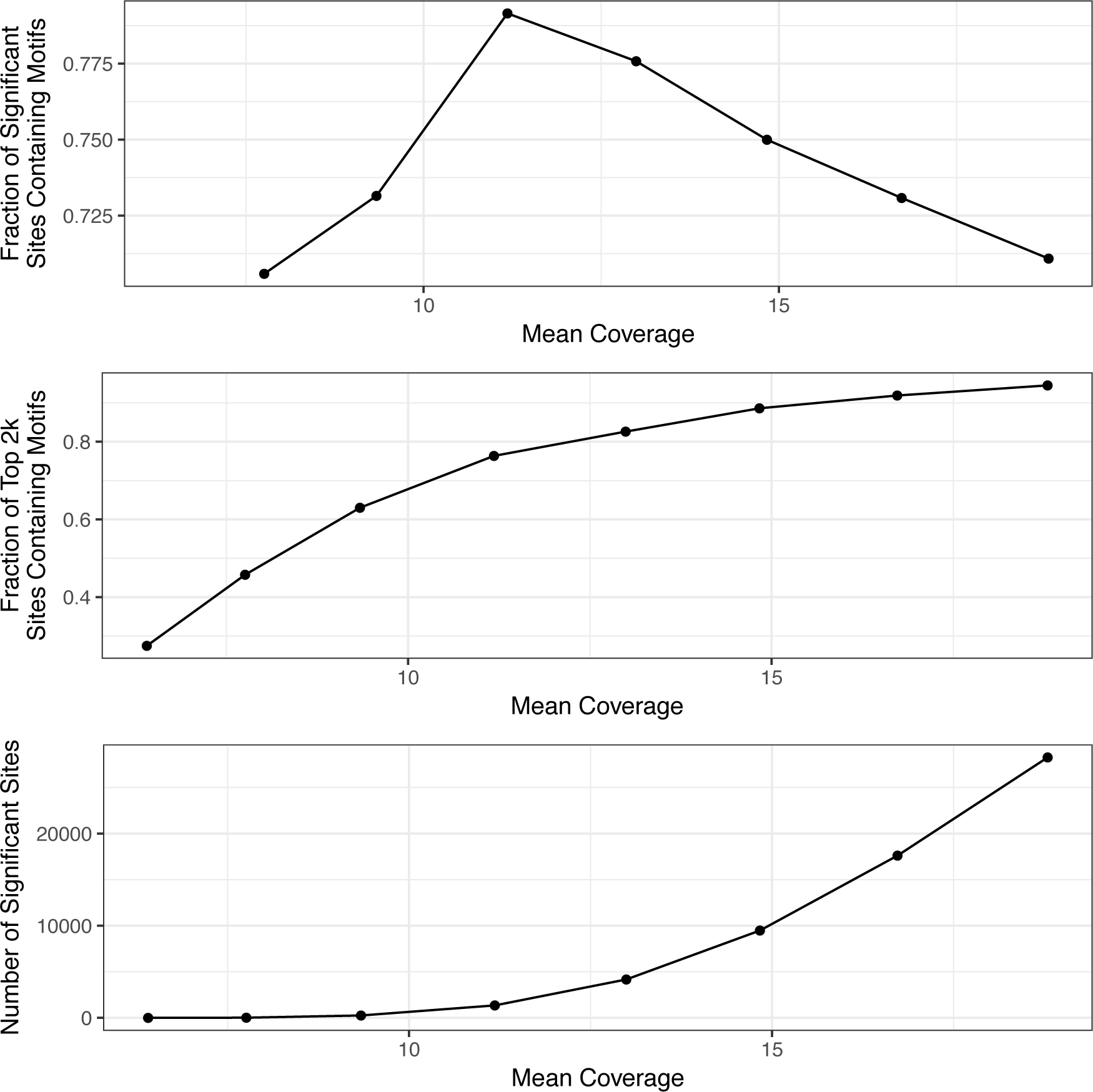
Relationship between statistical power (depth of coverage) and fraction of identified sites with known motifs in native E. coli samples over range of down-sampling to achieve different levels of strand-specific coverage (x-axis).

**Supplemental Figure 10.**
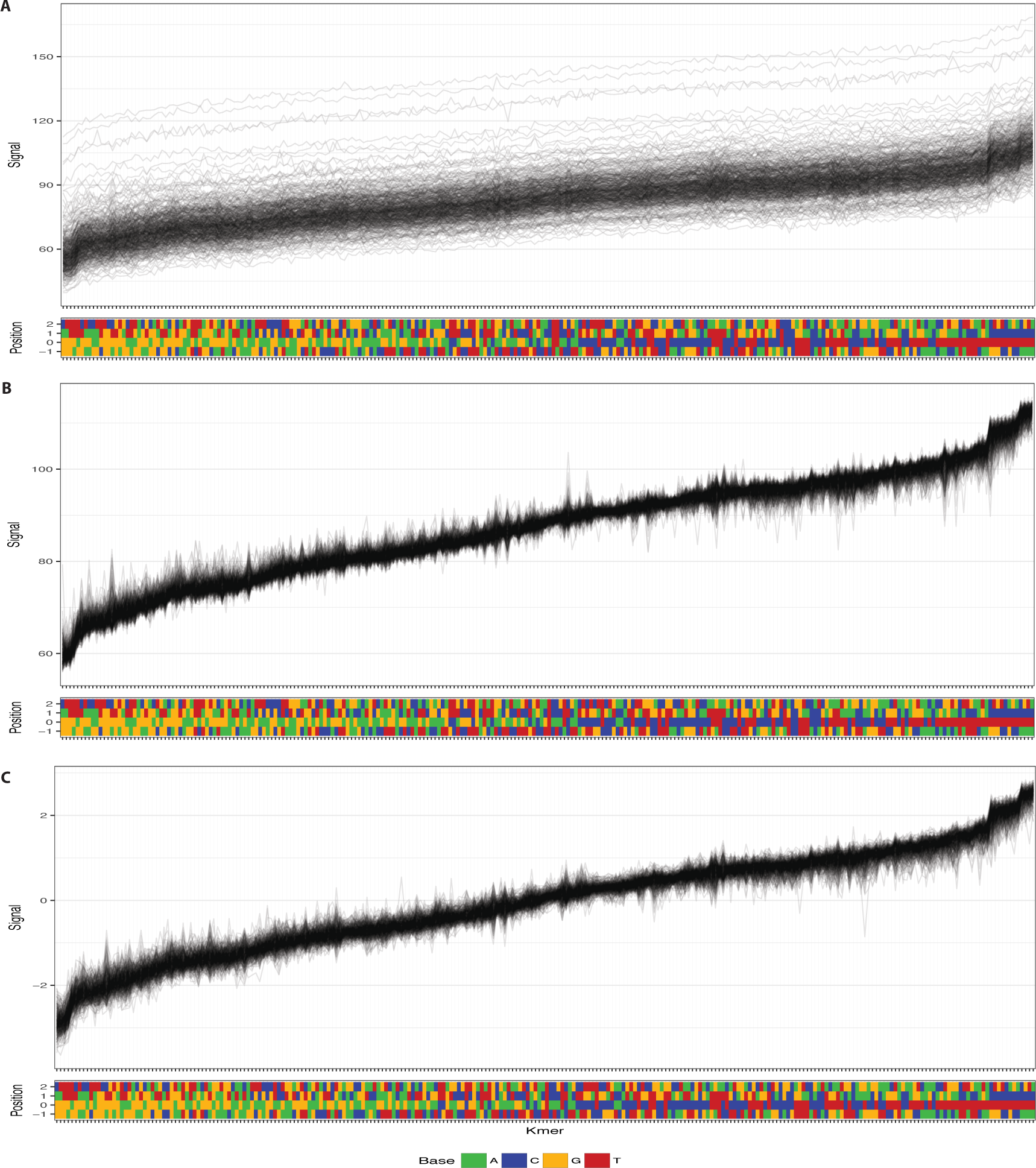
Oxford Nanopore Technologies raw pA normalization (A), corrected pA normalization (B) and median normalization (C). For each figure, left to right are 4-mers (from bottom to top position: one base already passed through the pore, the base at the center of the pore and two bases that have not yet passed though) ordered by mean signal across all reads. Each line represents the mean signal of one read across all 4-mers.

